# Design in Voxel Space Decode in SMILES Space: Plixer Generates Drug-Like Molecules for Protein Pockets

**DOI:** 10.1101/2025.07.15.664910

**Authors:** Jude Wells, Brooks Paige

## Abstract

We introduce Plixer, a two-stage generative model for de novo drug design that generates small-molecule ligand binders conditioned on an empty protein pocket. Plixer combines a conditional voxel inpainting network to generate 3D ligand hypotheses with an independently trained voxel-to-SMILES decoder that translates these voxel representations into valid chemical structures. By de-coupling the learning of spatial protein–ligand interactions from the learning of chemical grammar, our approach leverages large libraries of 3D ligand conformers to augment the limited data available for protein–ligand complexes. We show that this approach generates molecules with higher predicted binding affinity than recent methods.

## 1. Introduction

Discovering drug-like molecules that can effectively target specific protein pockets remains a central challenge in computational drug discovery. Traditional methods rely on either docking, which is computationally intensive, or on ligand-based screening, which is often limited by the availability of known actives. Recently, generative models have emerged as promising tools for exploring the vast chemical space in a data-driven manner. However, most of these generative models restrict themselves to training on the relatively few examples of experimentally resolved protein-ligand complexes.

In this work, we propose Plixer, a novel framework that integrates two complementary stages: (1) a conditional voxel inpainting model that generates a soft 3D voxel representation of a bound ligand given an empty protein pocket, and (2) a voxel-to-SMILES decoder that translates these voxelized ligand hypotheses into valid SMILES strings. The key innovation in Plixer is the decoupling of the tasks of learning spatial relations between protein pockets and ligands from the task of learning the chemical syntax and valid molecular structures. While only a limited dataset of protein–ligand complexes is available (a few thousand non-redundant samples), we can leverage millions of ligand-only examples to robustly train the decoder, thereby learning the manifold of valid molecules and energy landscape of 3D poses.

The inpainting model learns to predict the missing ligand voxels in the given pocket, while the decoder ensures that the inferred ligand lies on the manifold of chemically valid molecules. This two-stage framework not only improves the overall validity and diversity of generated SMILES strings but also allows for effective ranking of any candidate molecule via the conditional likelihood of the SMILES string.

## 2. Related work

### 2.1 Protein-Conditioned Generative Models for Ligand Design

Recent pocket-conditioned generators fall into three representational classes: (i) **voxel models** that predict an atomic-density grid inside the binding pocket, (ii) **graph models** that place discrete atoms via coordinates, and (iii) **SMILES models** that treat ligand design as a language problem. Voxel outputs can be converted to graphs by atom-fitting, and any graph can be rendered as a SMILES string, linking the three families. Orthogonally, their generation strategies can be *one-shot, autoregressive*, or based on *diffusion*.

**Voxel, one-shot**. The early voxel ligand predictor of Skalic et al. (2019) produced channel-wise densities but no discrete molecule. liGAN (Ragoza et al., 2022) and VoxBind (Pinheiro et al., 2024) improved this by decoding the density to a 3D graph with a fixed atom-placement heuristic, yet they cannot assign a likelihood to an arbitrary candidate and cannot exploit large ligand-only corpora during training. **Graph, autoregressive**. Pocket2Mol (Peng et al., 2022) and GraphBP (Liu et al., 2022) sequentially add atoms conditioned on the pocket. **Graph, diffusion**. DiffSBDD (Schneuing et al., 2024), TargetDiff (Guan et al., 2023), DiffBP (Lin et al., 2025a), ShapeMol (Chen et al., 2023), and PMDM (Huang et al., 2024) replace autoregression with a diffusion process that adds and moves atoms in 3D space.

#### Sequence-conditioned SMILES

DrugGPT (Li et al., 2023) and DrugGen (Sheikholeslami et al., 2025) use autoregressive language models to generate SMILES strings conditioned on a protein sequence.

#### Plixer

Our method is the first to generate a ligand voxel density and then translate it to SMILES with an autoregressive decoder trained on millions of 3D ligand structures. This design (i) yields a calibrated log-likelihood for any SMILES, enabling ranking as well as generation; (ii) leverages vast ligand-only data; (iii) allows us to sample multiple SMILES strings from a given voxel hypothesis. To assess Plixer as a ranking model that recovers hits from large libraries, we compare it against the latest SMILES string autoregressive model, DrugGen, (Sheikholeslami et al., 2025) and find that Plixer likelihood scores are more effective at separating hits from decoys although both machine learning methods are slightly worse than docking scores. For assessment of de novo binder generation, we compare against a recent autoregressive graph generative model, Pocket2Mol (Peng et al., 2022) that was shown to be competitive in a recent benchmark evaluation of many methods (Lin et al., 2025b). Here we find Plixer molecules are similar or better in computational assessment of target specificity.

## 3. Motivation

Our goal is to generate a drug-like molecule for a given receptor pocket *R*, i.e. to learn the map

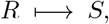

|

where *S* is a SMILES string. Directly fitting *P* (*S* |*R*) to the ≲ 10^4^ protein–ligand complexes in the PDB is challenging as the corpus is far too small to cover the vast chemical and syntactic space of valid SMILES. We therefore split the problem.

1. **Voxel inpainting**. A 3D convolutional U-Net deterministically produces a ligand atomic density grid *V* from the receptor atomic density grid *R*

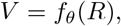

exploiting the shared spatial frame of receptor and ligand.
2. **SMILES decoding**. A vision–language model learns the *probabilistic* decoder

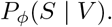

Because billions of drug-like molecules can be paired with their own voxelised conformers, *P*_*ϕ*_ is first pretrained on (*V, S*) pairs from the Zinc database of drug-like molecules (Irwin & Shoichet, 2005), then lightly fine-tuned on the smaller set of grids emitted by *f*_*θ*_.

Sampling from *P*_*ϕ*_ yields multiple chemically valid SMILES for a single voxel hypothesis, accommodating pockets that admit diverse chemotypes. Thus we harness the chemical knowledge of massive small-molecule corpora while exploiting 3D locality in *f*_*θ*_, making data-limited pocket-specific design tractable.

## 4. Methods

Plixer consists of two independent models that operate sequentially. The first component, a 3D voxel inpainting network, generates ligand-atom density grids conditioned on voxelized protein pockets. The second component translates these voxelized ligand representations into valid SMILES strings.

### 4.1 Protein-to-Ligand Voxel Generation (PocVox2MolVox)

The PocVox2MolVox model employs a 3D U-Net architecture to generate voxelized ligand representations from protein pocket inputs. The model takes as input a 4-channel voxel grid representing protein atom types (carbon, oxygen, nitrogen, and sulfur) and outputs a 9-channel voxel grid corresponding to ligand atom types (carbon, oxygen, nitrogen, sulfur, chlorine, fluorine, iodine, bromine, and other). We use a a cubic box of 24 Å with 0.75 Å voxels, resulting in a 32^3^ voxel grid. During training, we employ a combined loss function of equally weighted binary crossentropy (BCE) and Dice loss (Sudre et al., 2017). Early experiments with BCE alone typically produced all zero values while Dice loss alone resulted in false-positives on the rare-atom channels.

### 4.2 Voxel-to-SMILES Generation (Vox2Smiles)

The Vox2Smiles model follows an encoder-decoder architecture that combines a Vision Transformer (ViT) encoder with a GPT-style decoder to translate 3D voxelized ligands into SMILES strings. The encoder processes the 9-channel ligand voxel grid by dividing it into patches of size 4 × 4 × 4, resulting in a sequence of embedded patches. The ViT encoder consists of 8 transformer layers with 8 attention heads each. The decoder follows a GPT-2 architecture that autoregressively generates SMILES tokens conditioned on the encoded voxel representation. We use a custom SMILES tokenizer with a vocabulary size of 76 (one token for each of the common atom-types represented in Zinc plus additional tokens for the SMILES syntax plus special tokens).

### 4.3 Data Processing and Voxelization

Both protein pockets and ligands are represented as 3D voxel grids. Atoms are mapped to their respective channels based on atom type (e.g., carbon, oxygen, nitrogen). Each atom contributes to the voxel grid using a Gaussian distribution centered at the atom’s coordinates with the occupancy/decay rate based on the atom type’s van der Waals radius. During training, we apply random rotations and translations of up to 6 Å to ensure model robustness to different orientations and translations, including those induced by pocket mis-specification.

## 5 Results

### 5.1 Plixer can generate drug-like molecules that are target specific

We evaluate Plixer by measuring the following properties of the generated molecules: (i) the Vina docking score, (ii) hit-similarity enrichment, (iii) drug-likeness (QED), (iv) solubility: defined as the proportion of generated molecules in the desirable range for LogP (between -0.4 and 5.6) and (v) diversity: the number of unique molecules when generating one-per-target divided by the total number of targets (table 1). We find that Plixer performs better than the SMILES generative model DrugGen on all metrics and outperforms DrugGen and Pocket2Mol on predicted affinity metrics such as the Vina score and similarity enrichment, while Pocket2Mol outperforms Plixer on QED, LogP and diversity.

**Table 1.**
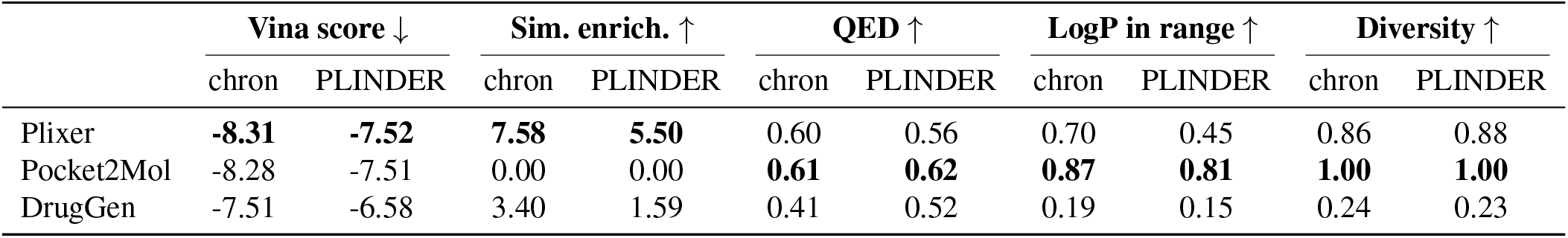
Docking and generative metrics on chronological (chron) and PLINDER-cluster splits of the PDB (best **in bold**).

|  | Vina score ↓ |  | Sim. enrich. ↑ |  | QED ↑ |  | LogP in range ↑ |  | Diversity ↑ |  |
| --- | --- | --- | --- | --- | --- | --- | --- | --- | --- | --- |
|  | chron | PLINDER | chron | PLINDER | chron | PLINDER | chron | PLINDER | chron | PLINDER |
| Plixer | **-8.31** | **-7.52** | **7.58** | **5.50** | 0.60 | 0.56 | 0.70 | 0.45 | 0.86 | 0.88 |
| Pocket2Mol | -8.28 | -7.51 | 0.00 | 0.00 | **0.61** | **0.62** | **0.87** | **0.81** | **1.00** | **1.00** |
| DrugGen | -7.51 | -6.58 | 3.40 | 1.59 | 0.41 | 0.52 | 0.19 | 0.15 | 0.24 | 0.23 |

### 5.2 Plixer can rank molecules by likelihood to recover hits

In addition to assessing properties of Plixer’s generated compounds, we can also use model to rank arbitrary compounds *S* against a target *R* according to Plixer’s SMILES likelihood score *P*_*ϕ*_ *(S* |*f*_*θ*_(*R*)). Given a set of compounds, the ROC-AUC score is computed from the model likelihoods and the binary labels for hit or decoy. The score has an intuitive interpretation as the expected probability that the model will rank a hit higher than a decoy. In our case, the hit is the single experimentally confirmed binder from the PDB file, while the decoy set is composed of all other molecules from the test set, excluding the confirmed binder. This ability to rank arbitrary molecules is not available in most generative models; as such, we only compare against DrugGen, and find Plixer outperforms this method, achieving a ROC AUC score of 0.58 on the PLINDER test set while using AutoDock Vina’s docking score to rank the same compounds achieves a ROC AUC score of 0.62 (table 5.2).

**Table 2.**
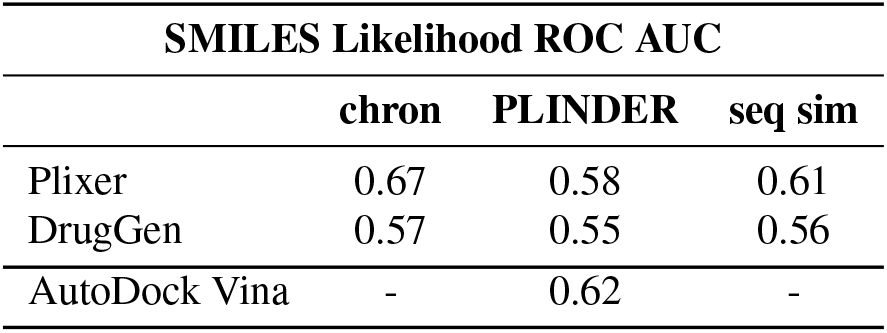
SMILES Likelihood ROC AUC scores on chronological, PLINDER and sequence similarity test splits

|  | SMILES Likelihood ROC AUC |  |  |
| --- | --- | --- | --- |
|  | chron | PLINDER | seq sim |
| Plixer | 0.67 | 0.58 | 0.61 |
| DrugGen | 0.57 | 0.55 | 0.56 |
| AutoDock Vina | - | 0.62 | - |

### 5.3 Structural similarity enrichment

In the context of drug discovery, methods are typically evaluated by considering the enrichment factor, defined as the ratio of the hit-rate observed among model-selected compounds to the hit-rate expected at random. Our evaluation in this paper is limited by the fact that we do not have experimental measurements for Plixer-generated molecules, therefore as a weak proxy for hit-identification we accept a generated molecule as a hit if it has a Morgan Fingerprint Tanimoto Similarity (MFTS) greater than 0.3 with the PDB experimentally confirmed binder. Thus, we define the similarity enrichment factor as the proportion of similarity hits among the generated molecules divided by the proportion of similarity hits among the decoy molecules. Where the decoy set is taken to be all molecules in the test set, excluding the true binder from the PDB. Table 1 shows Plixer is better than other benchmarked models at generating hits according to this metric, increasing the hit rate by a factor of 5.5 on novel protein pockets. Figure 3 shows the distribution of MFTS scores for Plixer-generated molecules with MFTS scores of decoys shown for reference.

**Figure 1.**
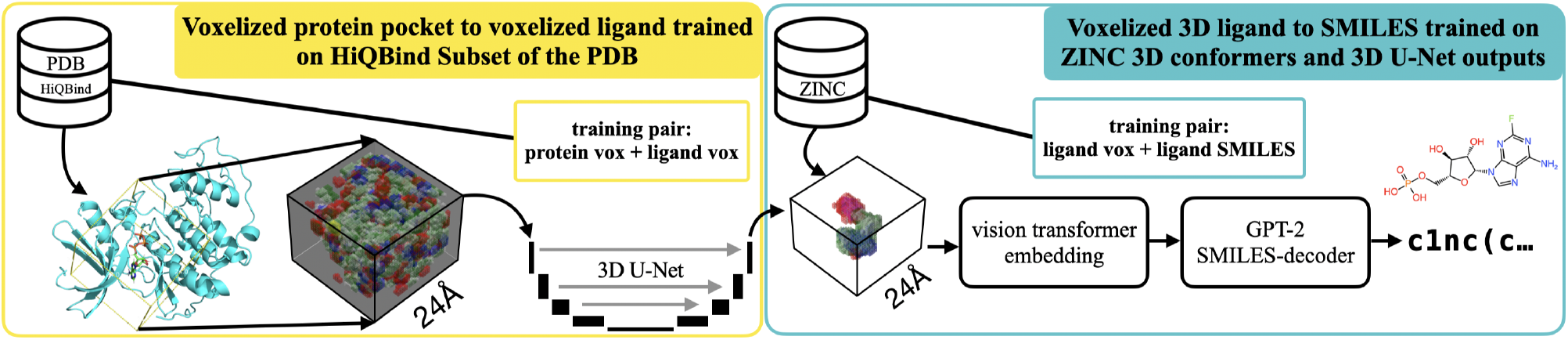
Plixer overview

**Figure 2.**
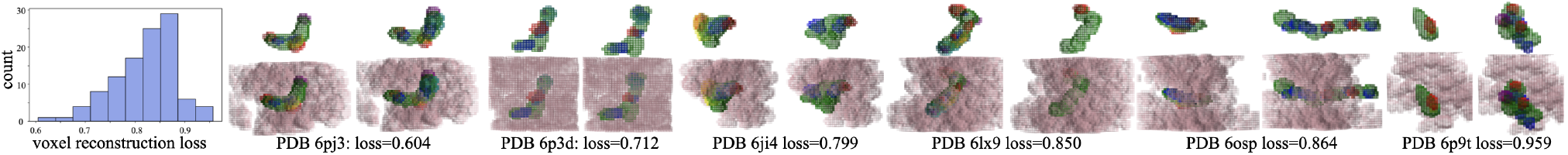
Distribution of voxel reconstruction loss scores on the PLINDER test set, with visualisations across different loss values with the Plixer prediction on the left and ground-truth on the right, lower row shows protein voxels in mauve.

**Figure 3.**
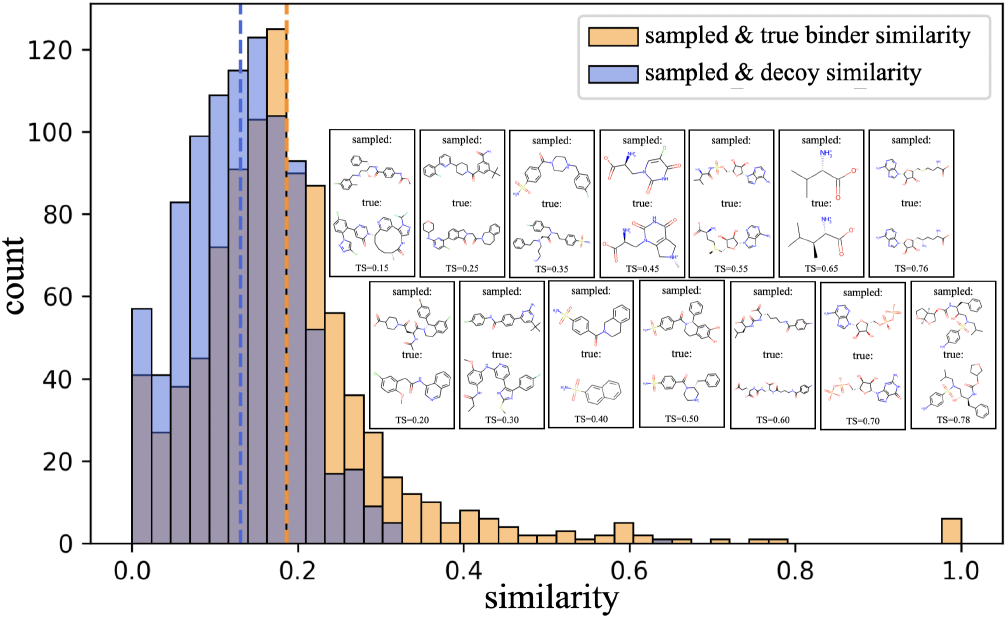
MFTS scores between the Plixer sampled molecule and the bound ligand from PDB versus decoys (a random PDB ligand from test set) and bound ligand from PDB. Additionally, we show paired molecules for a range of similarity scores with the Plixer sampled molecule on top and the true-binder PDB ligand on the bottom.

## 6. Discussion

Plixer shows that a seemingly modest recipe of CNNs on voxel grids, one-shot generation, and extensive data augmentation remains competitive with contemporary favourites such as GNNs with SE(3) equivariance and diffusion samplers. The chief obstacle for all pocket-conditioned generators is not model capacity but the limited availability of non-redundant protein–ligand complexes. By decoupling spatial reasoning (voxel inpainting) from chemical validity (voxel-to-SMILES decoding) and pre-training the decoder on millions of ligand-only conformers, we match or exceed state-of-the-art pocket-conditioned models on docking score and enrichment metrics. It is reassuring to see that model likelihoods on SMILES strings are discriminative in preferring true binders over decoys. Although Plixer likelihood scores are slightly less effective in ranking than docking scores from AutoDock Vina, the computation time is much faster.

### 6.1. Model limitations and potential improvements

#### Interpretability

Soft voxel densities are less interpretable than discrete atoms with coordinates; Adding an explicit coordinate prediction step would improve the usefulness of the model as a design tool. **One-shot artefacts**. The deterministic voxel generator can average multiple plausible chemotypes, sometimes producing non-physical densities. Autoregressive or diffusion generation may be preferable to mitigate this issue. **Feature representation**. Replacing atom-type channels with broader pharmacophore-like features (e.g. hydrogen-bond donors/acceptors) could enhance generalisation and novelty. **Protein flexibility**. The current model assumes a rigid receptor. Training on apo structures or on ensembles of receptor conformations could allow the network to account for pocket dynamics. **Non-structural binding data**. Many binders are confirmed via chemical assays without known complex structures. Incorporating such data via preference optimization or contrastive learning could enhance model performance.

## Code Availability

https://github.com/judeWells/plixer

## Impact Statement

Plixer aims to make pocket-guided molecule generation more efficient, potentially accelerating therapeutic discovery. As with any molecular design tool, dual-use risks exist but the principal barriers to causing deliberate or accidental harm are not significantly altered by this work.

## A Datasets

We trained our models on multiple datasets:

### A.1 Protein-Ligand training data

The PocVox2MolVox model was trained on the HiQBind dataset (Wang et al., 2025), a filtered collection of protein-ligand complexes derived from the PDB. We first applied a chronological split keeping all entries from 2020 and later for the test set with the remainder held for training and validation. The validation dataset was created by applying MMSEQS2 (Steinegger & Söding, 2017) to generate clusters with 30% sequence identity and at least 50% coverage. 10% of the protein clusters are removed from training to be used as validation samples. Ligands were also clustered using Morgan fingerprint representation and the rdkit Butina clustering algorithm each sample in the dataset gets a cluster identity which is formed from the tuple of the protein cluster and the ligand cluster and one training epoch is defined as a single pass over all clusters. A chronological split of the PDB is insufficient to determine model performance on novel proteins, therefore we created two subsets of the chronological test set, one using the PLINDER (Durairaj et al., 2024) pocket clusters (removing all samples from the test set which shared a PLINDER pocket cluster label with any training sample) and one using MMSEQS protein sequence similarity: removing all samples from the test set which had 30% sequence similarity with any protein in the training set.

### A.2 Ligand-only training data

ZINC20: The Vox2Smiles model was initially trained on a subset of the ZINC20 database (Irwin et al., 2020), which provides 3D conformers of small molecules. This enabled the model to learn the relationship between 3D structural representations and SMILES strings without requiring protein context. Combined Dataset: For fine-tuning the Vox2Smiles model, we created a combined dataset that includes both ZINC20 molecules and outputs from the PocVox2MolVox model. 15% of the training samples came from PocVox2MolVox outputs while 85% came from ZINC20. Additionally, we filtered PocVox2MolVox outputs to include only those with a Dice+BCE loss value below 0.7 to ensure quality voxel representations.

## B Performance at different levels of protein novelty

Generative models for structural biology have been found to generalise poorly beyond their training data (Buttenschoen et al., 2024). We observed moderate decreases in our model performance when we assess on the more strict splits using PLINDER cluster exclusion or sequence similarity exclusion compared to the chronological split. However, figure 4 suggests that MFTS is not particularly correlated with the maximum sequence similarity found between the test example and another example in the training set. We attribute this to two factors: first, we use a strict validation split (not just chronological) and stop training when the validation loss starts to increase. Second, similar proteins in the PDB may have quite different ligands present in the pocket.

**Figure 4.**
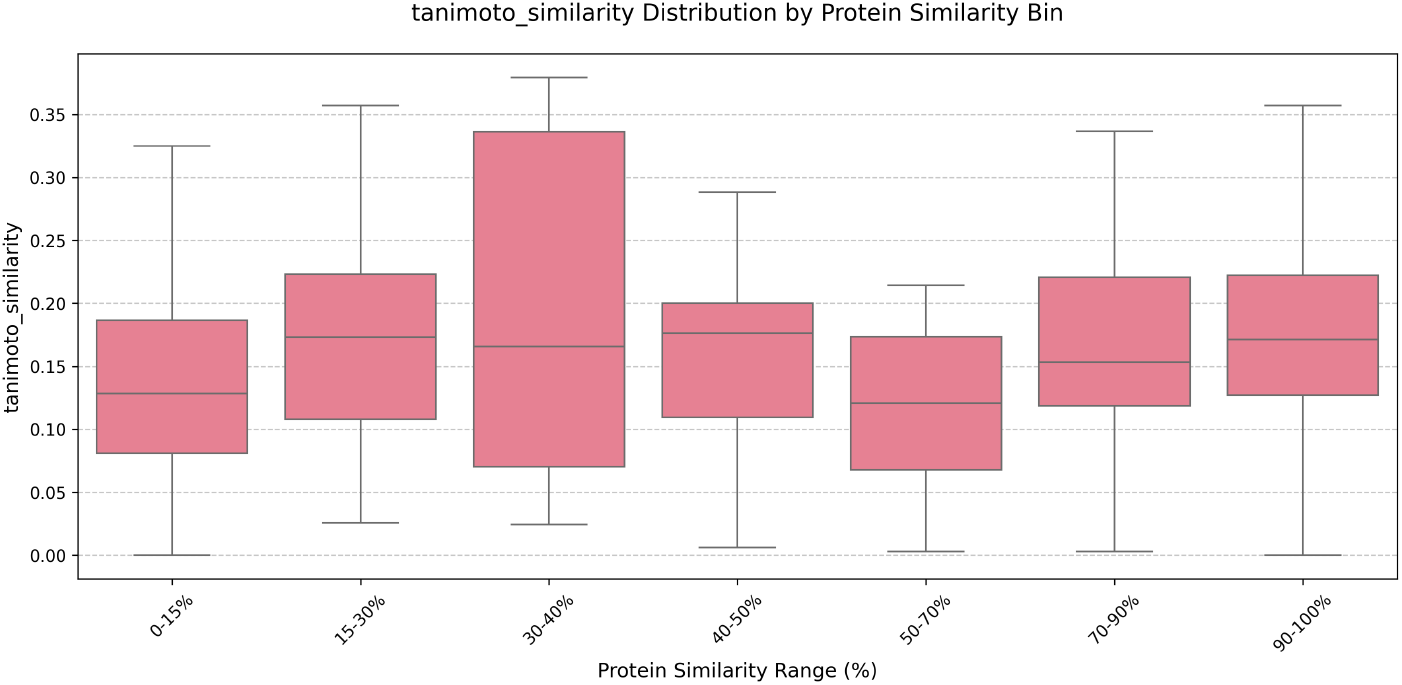
Morgan fingerprint tanimoto similarity (MFTS) of generated molecule and PDB molecule for different levels of maximum protein similarity with any sample in the training set

## Notes

https://github.com/judeWells/plixer

### Competing Interest Statement

The authors have declared no competing interest.

### Funding

- Wellcome Trust: 218529/ Z/19/Z

